# Identification and Characterization of novel long non-coding RNAs in vascular smooth cells

**DOI:** 10.1101/2023.01.06.523019

**Authors:** Charles U. Solomon, David G. McVey, Catherine Andreadi, Peng Gong, Lanka Turner, Sonja Khemiri, Julie C. Chamberlain, Tom R. Webb, Nilesh J. Samani, Shu Ye

## Abstract

A significant portion of the RNA produced from the human genome consists of long non-coding RNAs (lncRNAs). These molecules tend to have lower levels of expression, are more specific to certain tissues, and show greater variation in expression between individuals compared to protein-coding messenger RNAs (mRNAs). LncRNAs have been linked with regulatory roles in gene expression and genome architecture. There is growing evidence that lncRNAs play important roles in many biological processes and diseases, and a number of lncRNAs have been identified as potential therapeutic targets. Here, we report the identification and characterization of the lncRNA landscape of vascular smooth muscle cells (VSMC). We used an ensemble of bioinformatics tools to identify 329 novel lncRNAs from a large VSMC RNA-Seq dataset. We found that majority of the novel lncRNAs are natural antisense transcripts of protein-coding genes. In addition, we predicted cellular localization and potential miRNAs that targets the novel lncRNAs and found that most localize in the cytoplasm and that miRNA target site ranged from 2-889 sites on each novel lncRNA. Furthermore, we identified co-expressed lncRNAs that correlate with the proliferation, migration and apoptosis of vascular smooth muscle cells. These results suggest that we have identified a diverse set of previously unknown lncRNAs that may be involved in important regulatory pathways in vascular smooth muscle cells.

## Introduction

Long non-coding RNAs (lncRNAs) constitute a large proportion of the RNA pool transcribed from the human genome.^1^ They generally have lower expression levels, greater tissue-specificity in expression, and greater variability in expression between individuals, than protein-coding mRNAs.^2-6^. There is growing evidence that lncRNAs play important roles in many biological processes and diseases,^7^ and a number of lncRNAs have been identified as potential therapeutic targets.^8-10^ However, there is still a lack of complete annotation of lncRNAs in human and other species^11^ and our current understanding of their expression patterns and biological roles is still rather incomplete.

Whole-genome RNA-sequencing (RNA-Seq) provides a means for identifying known and novel lncRNAs (as well as other types of RNA). RNA-Seq of a number of different types of tissue has been conducted by the GTEx Consortium and other investigators, and data from these efforts have helped in gaining important biological insights on lncRNAs.^12^ However, since tissue samples generally consist of a mix of different types of cell and since the expression of some lncRNAs can be cell-type specific,^13^ RNA-Seq of a single type of cell may provide a valuable alternative approach for precise characterization of the lncRNA transcriptome of a given cell type. This could be combined with phenotypic assays for the investigation of relationships between lncRNA expression and cellular behavior.

Vascular smooth muscle cells (VSMCs) are a major cell type in arteries and play major roles in vascular diseases, e.g. changes in VSMC proliferation, migration and apoptosis are key processes in the pathogenesis of atherosclerosis, a common pathological condition that underlies ischemic heart disease and contributes to the development of hypertension, aneurysm and stroke.^14^ Several lncRNAs have been shown to modulate VSMC functional characteristics.^15, 16^ However, it is plausible that there are still other lncRNAs that can also influence VSMC behavior, which are yet to be identified.

In this study, we performed whole-genome RNA-Seq and behavior assays of a large collection of VSMCs from different individuals (n=1,499), and used this large dataset to characterize the VSMC lncRNA transcriptome and to systematically investigate relationships of the expression of different lncRNAs with VSMC proliferation, migration and apoptosis, respectively.

## Methods

### VSMC isolation and storage

VSMCs were isolated from the artery of umbilical cords from 2016 donors, using a reported method^17^. Aliquots of VSMCs of passage 3 were either stored in freezing medium under liquid nitrogen or in RNAlater solution (Sigma) at -20°C. Umbilical cord tissues used in this study were collected and provided by the Anthony Nolan Trust with ethical approval and informed consent of donor’s parents.

### VSMC behavior assays

#### Proliferation assay

Cells were seeded at a density of 5,000 cells per well in 0.2% gelatin-coated 96-well plates. After overnight incubation, the cells were incubated with 10µM EdU for a period of 6 hours. After this incubation, the cells were fixed with 3.7% formaldehyde for 15 minutes. EdU incorporation was detected using the BaseClick EdU HTS 488 kit (Sigma, BCK-HTS488-20) using the fluorophore 6-FAM, following the manufacturer’s protocol. Cells were then incubated with 1.6µM Hoechst 33342, and imaged on an Operetta CLS High-Content Analysis System (Perkin Elmer) with a 10x air objective and 21 fields per well. Following imaging, image analysis using Harmony 4.8 was performed. Firstly, the nuclei were segmented using the Hoechst 33342 channel followed by calculation of the intensity of 6-FAM staining. Cells with maximum 6-FAM staining above a threshold of 2000 were classified as EdU-positive, whilst cells with maximum intensity below 2000 were classified as EdU-negative. The EdU-positive cells were further segregated into “high” and “low” categories based on intensity. Cells with maximum intensity values >10,000 were classified as “high” whilst those with maximum intensity between 2000 and 10,000 were classified as low. The percentage of total cells for each category was calculated.

#### Migration assay

Cells were seeded at a density of 1,500 cells per well in 0.2% gelatin-coated 96-well plates. After incubation overnight, the cytoplasm of the cells was stained with 10µM CellTracker Green CMFDA (ThermoFisher) following the manufacturer’s protocol, whilst the nuclei were stained with 0.4µM Hoechst 33342. After staining, the cells were incubated in media containing 2% foetal calf serum and 0.4µM Hoechst 33342. The cells were then imaged every 1 hour for 16 hours using an Operetta CLS High-Content Analysis System (Perkin Elmer) with a 10x air objective and 21 fields per well. Image analysis was performed to segment the cell nuclei and cytoplasm, followed by individual cell tracking over the course of the assay. Cells fully tracked for the full 16 hours were used to calculate the straightness, speed, accumulated distance, and displacement parameters using Harmony 4.8 (all cells with 2 or more consecutive time points were also used to calculate the migration speed).

#### Apoptosis assay

We performed apoptosis assay on VSMCs from 1861 donors. Cells at passage 3 were seeded into 96-well plates (CellCarrier Ultra, Perkin Elmer) coated with 0.2% gelatin at a density of 800 cells per well, with 4 replicate wells for cells from each donor. After overnight incubation, the cells were stained with Hoechst 33342 (1.6µM) and propidium iodide (0.5µM), the latter of which only stains membrane-compromised cells, and imaged on an Operetta CLS High-Content Analysis System (Perkin Elmer) with a 10× air objective and 9 imaging fields. The plate was removed from the instrument and the media replaced with media containing Hoechst 33342, propidium iodide and the apoptosis inducer staurosporine (2.5µM). The plate was returned to the Operetta CLS and incubated for 30 minutes before imaging the plate every 30 minutes for 16 hours. Upon completion of the assay, the images were analyzed using Harmony 4.8 software (Perkin Elmer). The cell nuclei were first segmented using the Hoechst 33342 channel and nuclear morphology and intensity parameters were calculated. The area of the nucleus was compared between the initial image and the 30-and 60-minute timepoints. Also, nuclear fragmentation was assessed using the fragmentation index (the coefficient of variation in nuclear Hoechst 33342 pixel intensity), and comparisons between the initial, pre-staurosporine treatment images and the fragmentation index at 30 and 60 minutes post-staurosporine treatment were made. Cells were also assessed for propidium iodide staining; those with positive staining resulting from compromised cell membrane integrity were classified as “dead”. The percentage of cells classified as “dead” at each time point were calculated and also used to determine the time taken for 50% of the cells to become propidium iodide staining positive.

### RNA isolation, strand-specific RNA-Seq library preparation and sequencing

Total RNA was extracted from an aliquot of passage-3 VSMCs in RNAlater solution, with the use of the Biobasic EZ-10 DNAaway RNA miniprep kit (Biobasic), according to the manufacturer’s protocol with additional wash steps to ensure complete removal of residual salts from the samples. RNA concentration and integrity was assessed by RNA BR and RNA IQ assays using a Qubit4 instrument (ThermoFisher). Samples were used for RNA-sequencing if they had a concentration >70ng/µl and an RNA integrity number (RIN) ≥ 6.8. Analysis of the RNA samples using a NanoDrop 8000 was also performed, with a 260:280 and 260:230 acceptance threshold of >=2.

A strand specific library with rRNA removal was prepared from total RNA of each VSMC sample, and 150bp paired-end sequencing at a 30 million read depth was performed using the Illumina platform, carried out by Novogene.

### RNA-Seq data quality control and mapping

Read quality was assessed with FastQC v0.11.5 (http://www.bioinformatics.babraham.ac.uk/projects/fastqc). Adapters were trimmed with BBMap v38.51 (https://www.osti.gov/biblio/1241166). STAR v2.7.1a^18^ was used to map reads to the human reference genome file, Homo_sapiens.GRCh38.dna_sm.primary_assembly.fa and annotation file Homo_sapiens.GRCh38.100.gtf, both downloaded from Ensembl^19^ (Accessed 10/06/2020). The non-default STAR options used for read mapping were –twopassMode Basic, –outSAMunmapped Within, and –limitSjdbInsertNsj 2000000.

### Novel lncRNA identification pipeline

We implemented a pipeline outlined in Supplementary Figure I and briefly described below, to identify novel IncRNAs.

#### Transcript assembly and detection of candidate novel lncRNA

Mapped reads were assembled and merged with Stringtie v2.1.1.^20^ lncRNA annotation from LNCipedia^21^, lncipedia_5_2_hg38.gtf (Accessed 03/08/2020) was combined with Homo_sapiens.GRCh38.100.gtf using cuffmerge v2.2.1^22^. The merged assembly was compared to the combined Ensembl-LNCipedia annotation using GffCompare v0.12.1^23^. Transcripts that were 200 nt in length, had more than two exons, and whose GffCompare classification codes were “i”, “u”, “x” were selected as candidate novel lncRNA. Transcript sequences were extracted from Homo_sapiens.GRCh38.dna_sm.primary_assembly.fa with GffRead v0.12.2^23^ and SeqKit^24^.

#### Filtration of candidate novel lncRNA

Sequences of candidate lncRNAs were assessed for coding potential with CPC^25^, CPAT^26^ and PLEK^27^ and those reported as non-coding were selected. Also, the candidate lncRNAs were filtered with FEELnc_filter.pl module with the option –monoex=-1 to remove spurious transcripts, and their coding potential assessed with FEELnc_codpot module with ensemble protein_coding and lncRNA transcripts serving as training datasets.^28^ The sequences of candidate lncRNAs were translated to all possible six frames and blasted against Uniprot^29^ and Pfam^30^ databases using blast+ v2.9.0^31^. Transcripts whose translated frames did not have any significant (e-value 10×^-10^, alignment length 10, and amino acids and identity 95%) blast hits were selected as non-coding. Transcripts that passed these six filtration steps and not currently annotated in Ensembl Homo_sapiens.GRCh38.100.gtf and lncipedia_5_2_hg38.gtf were considered novel lncRNAs.

#### Annotation of novel lncRNA

The novel lncRNAs were annotated with FEELnc_classifier.pl module^28^. Homo_sapiens.GRCh38.100.gtf served as reference annotation and the best matches were selected. Further information on each transcript was extracted from the transcript GTF file and added to the FEELnc classification using a custom R script.

### In silico functional characterization of novel lncRNA transcripts

#### Subcellular localization analysis of novel lncRNA transcripts

We predicted the subcellular localization of the novel lncRNA transcripts using two online tools; iLoc-lncRNA (Su et al 2018) (Accessed 18/03/21) and LoclncRNA (Wang et al 2021) (Accessed 18/03/21). iLoc-lncRNA predicts localization in four cell compartments; nucleus, cytoplasm, ribosome and exosome while LoclncRNA predicts localization in just nucleus and cytoplasm. To make the predictions from both tools comparable we classed ribosome and exosome predictions by iLoc-lncRNA as cytoplasm. We selected only transcripts with consensus predictions from both tools.

#### miRNA Target Prediction

We used Probability of Interaction by Target Accessibility (PITA) (Kertesz *et al*, 2007) to predict novel lncRNAs that are targets of miRNAs. Human mature miRNAs were downloaded from miRbase v22 (Kozomara *et al*, 2019) (accessed 04/06/2021). Default PITA settings were used to predict the miRNA targets and only sites with ΔΔG < 15 kcal/mol were considered.

### Gene expression quantification and normalization

We used kallisto^32^ to quantify gene expression. Transcript sequences of Stringtie merged reads were extracted from Homo_sapiens.GRCh38.dna_sm.primary_assembly.fa with GffRead, indexed and each RNASeq sample was quantified with kallisto using the following options -b 100 –rf-stranded. The R package tximport v1.16.1^33^ was used to obtain gene-level count of expression. DESeq2 v1.28.0^34^ was used to normalize the gene expression data.

### Correlation analysis

Spearman correlation coefficients were calculated using the R package psych (https://CRAN.R-project.org/package=psych).

### Weighted gene co-expression analysis (WGCNA)

Gene name and biotype of expressed genes were obtained with Biomartr v2.44.0^35^. We then extracted lncRNAs, including the novel lncRNAs we just discovered, from the gene expression data. The lncRNA expression data and VSMC phenotype data was used as input for WGCNA v1.69^36^. One sample was judged an outlier and removed based on sample clustering dendrogram. After removing lncRNAs with excessive missing values, 16256 lncRNAs were used for network analysis. We chose a power of 5 based on the scale free topology and mean connectivity plots (Supplementary Figure II and III). The network was constructed with the WGCNA function, blockwiseModules with the following arguments networkType = “signed”, corType = “bicor”, minModuleSize = 30, mergeCutHeight = 0.25. Module hub genes were obtained with the function chooseTopHubInEachModule. Pearson correlation between module eigengenes and VSMC phenotypes was calculated with the WGCNA function, cor and plotted with the function, labeledHeatmap.

### GO Term analysis of partner protein-coding genes of lncRNAs in different modules

Because most lncRNAs still lack functional annotation, we could not carry out gene ontology analysis directly with lncRNAs assigned to different WGCNA modules. As a proxy, we performed gene ontology analysis using protein-coding genes that are located nearby lncRNAs in the genome (in this paper, we refer to these nearby protein-coding genes as partners). For each module, partner protein-coding genes located within 10kb of the module’s lncRNAs were obtained with BEDTools v 2.25.0 (Quinlan and Hall, 2010). The VSMC expression profile of lncRNAs and their partner protein-coding genes were correlated. Partner protein-coding genes with r ≥ 4 (p-value ≤ 0.05) were used to perform gene ontology analysis with ToppGene (accessed 12/08/2021) (Chen et al 2019). Redundant gene ontology terms were filtered out with REVIGO (accessed 15/09/2021) (Supek et al. 2010) in order to make plots.

## Results

### Known and novel lncRNA detected in the VSMC transcriptome

In this study, we performed an RNA-Seq analysis on a large collection of vascular smooth muscle cells (VSMCs) derived from umbilical arteries of different donors (n=1,486). In brief, a strand specific library with rRNA removal was prepared for each sample, and 150bp paired-end sequencing at a 30 million read depth was performed using the Illumina platform. The RNA-Seq generated an average of 47,077,173 raw reads per sample, with an average of 43,496,830 reads after trimming and 93.07% of reads being uniquely mapped (Supplementary Table I).

From this RNA-Seq dataset, we detected transcripts from a total of 60,790 genes. Of these 60,790 genes, 32.8% were protein-coding genes, 28.1% were lncRNA genes, and the remainder were pseudogenes, genes of other types of RNA (Figure 1 and Supplementary Table II). As described elsewhere, a multi-dimensional scaling (MDS) analysis comparing the transcriptomes of our VSMC samples with reported transcriptomic data from human coronary artery smooth muscle cells^37^ and transcriptomic data from the GTEx Portal^13^ for other cell/tissue types showed that our VSMC samples were very similar to human coronary artery smooth muscle cells but dissimilar to other cell types.

**Figure 1:**
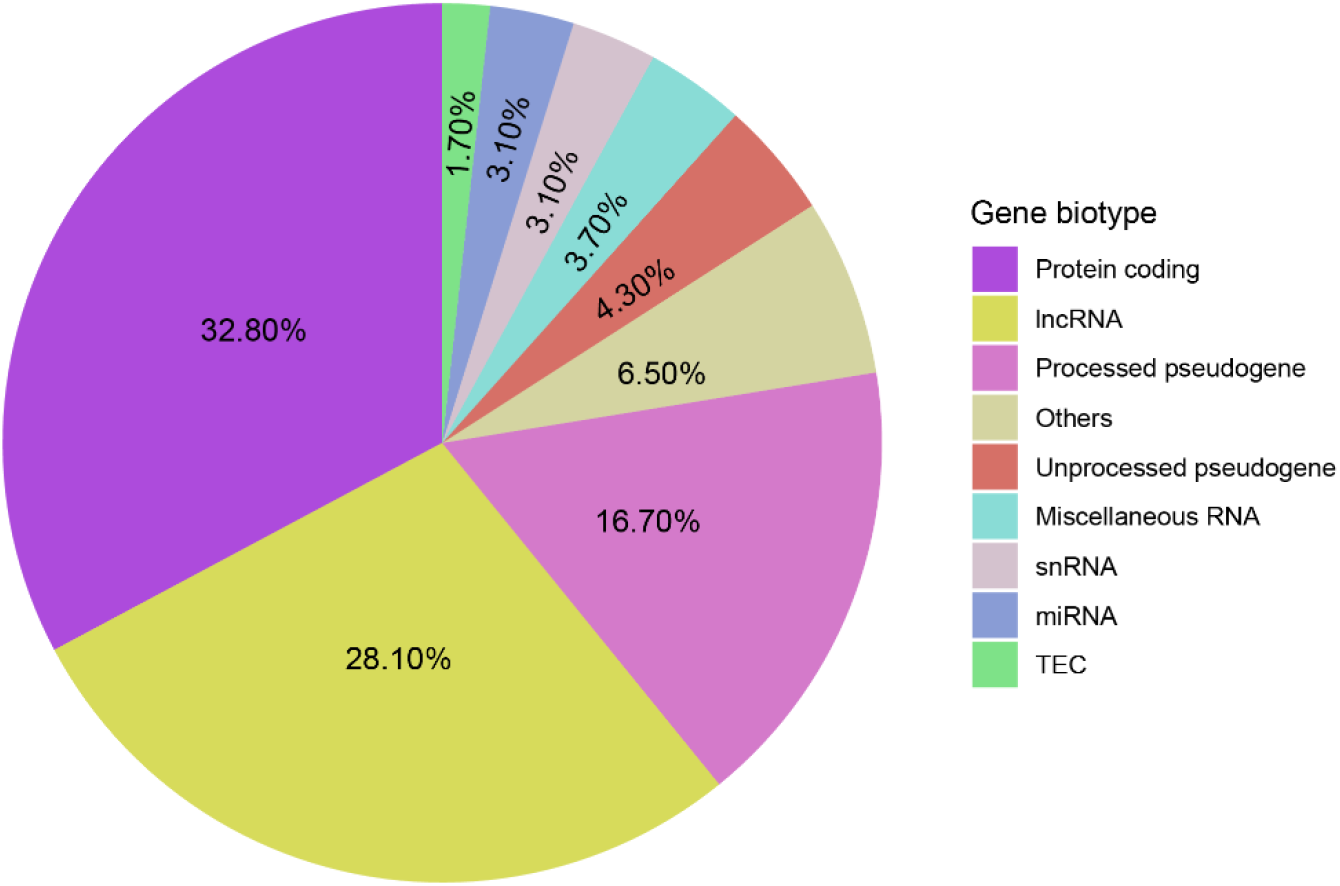
The percentages of different types of genes detected in VSMCs in this study. TEC; To be Experimentally Confirmed.

The use of a strand specific, directional library prepared from total RNA, with removal of the abundant rRNAs to enrich non-rRNA transcripts, for RNA-Seq in our study made our data high suitable for lncRNA discovery and analysis. We used an analytic pipeline described in Supplementary Figure I to identify novel lncRNAs. The pipeline included methods that had been previously used to identify lncRNAs^38-41^ as well as multiple well-established software to confirm that the lncRNAs identified had no strong coding potential and to verify that none of them represented a coding mRNA for any protein recorded in the Uniprot or Pfam databases.

This pipeline found 329 novel lncRNAs and additionally detected 244 previously annotated lncRNAs, in our VSMC RNA-Seq dataset (Supplementary Table III). Of the 329 novel lncRNAs, 278 were genic transcripts and the remaining 51 were intergenic, with the vast majority being antisense to their partner mRNA transcripts (Table 1). An analysis using FEELnc^28^ indicated that the novel lncRNA transcripts detected encoded a total of 185 genes.

**Table 1:** Classification of novel lncRNAs.

| Novel VSMC lncRNAs transcripts |  |  |  |  |  |  |  |  |
| --- | --- | --- | --- | --- | --- | --- | --- | --- |
| Intergenic (51) |  |  | Genic (278) |  |  |  |  |  |
| Same Strand | Convergent | Divergent | Containing (39) |  | Nested (135) |  | Overlapping (104) |  |
| 17 | 23 | 11 | S | AS | S | AS | S | AS |
|  |  |  | 14 | 25 | 9 | 126 | 4 | 100 |

S, Same sense; AS, antisense.

572Using *in silico* tools we predicted cellular localization and potential miRNAs that targets the novel lncRNAs. We obtained consensus localization predictions for 161 out of 329 novel lncRNAs using iLoc-lncRNA (Su et al 2018) and LoclncRNA (Wang et al 2021). Fifteen novel lncRNAs were predicted to localize in the nucleus while 146 were predicted to localize in the cytoplasm (Supplementary Table VI). The proportion of lncRNA predicted to localize in the cytoplasm and nucleus are similar to previous reports (Heesch et al 2014; Carlevaro-Fita and Johnson, 2019). In addition, the predicted number of miRNAs that target sites in individual novel lncRNAs ranged from 2 to 889 miRNAs per transcript (Supplementary Table VII).

### Associations between lncRNA expression and VSMC behaviour

To test associations between the expression of the various lncRNAs and VSMC behavior, we first carried out a Spearman correlation analysis of the expression value of each lncRNA in relation to results from proliferation, migration and apoptosis assays, respectively, on VSMCs from n=1,499 individuals in our VSMC bank. The analysis identified a total of 572 lncRNAs whose expression highly significantly correlated with VSMC behavioral parameters (significance threshold, correlation coefficients and P values are described in Supplementary Table IV). Some of these lncRNAs have previously been implicated in VSMC biology, e.g. ANRIL has been reported to promote VSMC migration^42^ and in agreement, its expression positively correlated with VSMC migration in our study (Figure 2A), whereas H19 has been described to inhibit VSMC apoptosis and our study observed an inverse association between H19 expression and VSMC apoptosis. Additionally, our study showed associations of many other lncRNAs with various VSMC behavior parameters (Supplementary Table IV).

**Figure 2:**
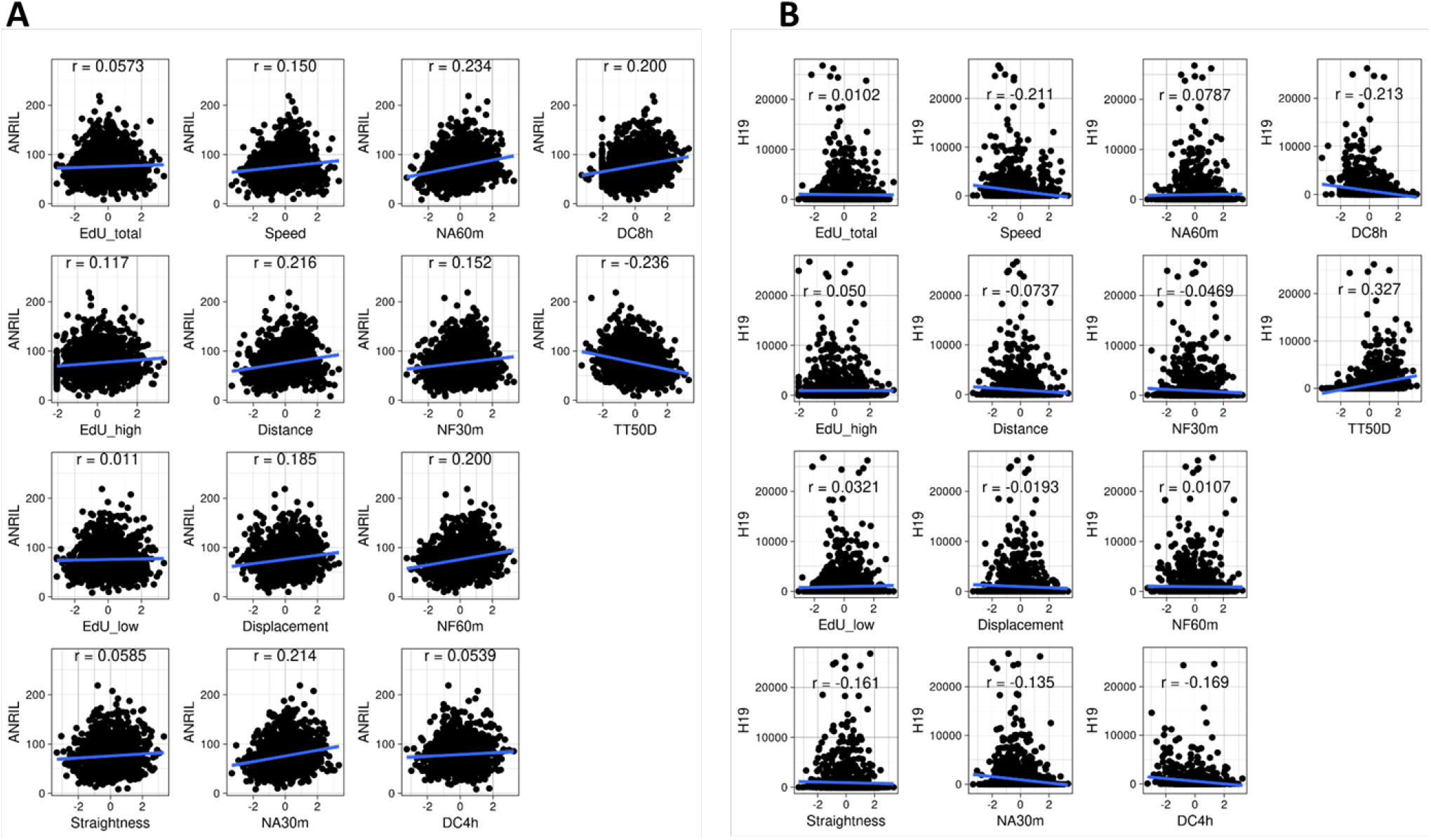
Correlations of ANRIL and H19 with VSMC behavior parameters.

As it is expected that VSMC behavior is influenced by many lncRNAs concurrently, we performed a weighted gene co-expression network analysis (WGCNA)^36^. The WGCNA algorithm clustered the lncRNAs detected in our VSMC samples into 6 co-expression modules, which were designated blue, brown, green, grey, turquoise, and yellow, respectively (Figure 3A, Table 2 and Supplementary Table V). The green module had the smallest number of genes, whilst the grey module consisted of the largest number of genes, noting that the WGCNA algorithm typically assigns genes whose expression has little variation to the grey module.

**Figure 3.**
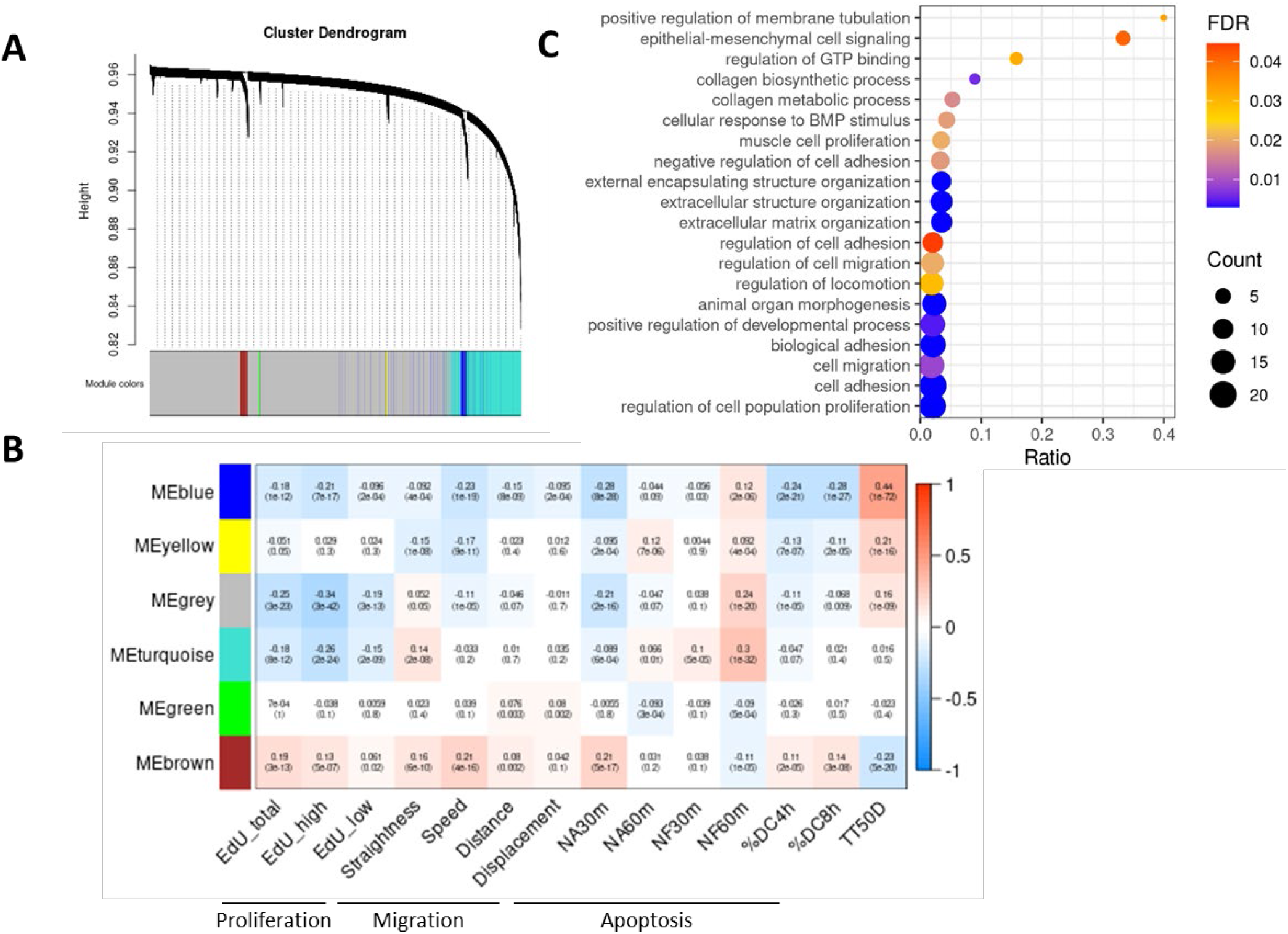
Associations of lncRNA gene modules with VSMC behavior. **A**. Clustering dendrogram of gene expression values based on topological overlap and module assignment from weighted gene co-expression network analysis (WGCNA) of VSMC RNA-sequencing data. **B**. Heatmap representation of correlations of gene modules with VSMC behavior parameters. Values shown are correlation coefficients and p-values (in brackets). The prefix ME in each module name stands for module eigengene. NA30m and NA60m: change in nuclear area at 30 and 60 minutes post treatment with the apoptosis inducer staurosporine; NF30m and NF60m: change in nuclear fragmentation index at 30 and 60 minutes post staurosporine-treatment; %DC4h and %DC8h: the percentage of dead cells (propidium iodide positive) at 4 and 8 hours post staurosporine-treatment; TT50D: the time in minutes for 50% of cells to become propidium iodide positive post staurosporine-treatment. **C**. Biological process category of blue module GO terms.

**Table 2:** WGCNA modules and hub genes of VSMC lncRNAs.

| Module | No. of genes | Hub gene | Symbol | Description |
| --- | --- | --- | --- | --- |
| Blue | 504 | ENSG00000229847 | EMX2OS | EMX2 opposite strand/antisense RNA |
| Brown | 336 | ENSG00000242125 | SNHG3 | small nucleolar RNA host gene 3 |
| Green | 47 | ENSG00000269243 | NA | NA |
| Grey | 12603 | NA | NA | NA |
| Turquoise | 2644 | ENSG00000241743 | NA | NA |
| Yellow | 122 | ENSG00000204387 | SNHG32 | small nucleolar RNA host gene 32 |

NA; not available

The WGCNA analysis revealed significant associations of the co-expressed gene modules with the various VSMC behavior parameters (Figure 3B). The strongest and statistically most significant association observed was an inverse correlation between the blue module and apoptosis, especially the parameter TT50D (the length of time before 50% of the VSMCs were found to have died after treatment with the apoptosis inducer staurosporine, *r* = 0.44, P = 1×10^−72^, Figure 3B).

To further assess the associations of gene modules with VSMC behavior, we used the WGCNA package to examine the correlation between module membership (correlation between lncRNA expression value and module membership eigengene value) and VSMC behavior association (correlation between lncRNA expression value and VSMC behavioural parameter value). Again, the blue module and TT50D showed the strongest correlation (r = 0.74, p = 1.7×10^−88^)(Figure 4), compared with the relationships of the other modules and VSMC phenotypes (Supplementary Figure IV-VI).

**Figure 4.**
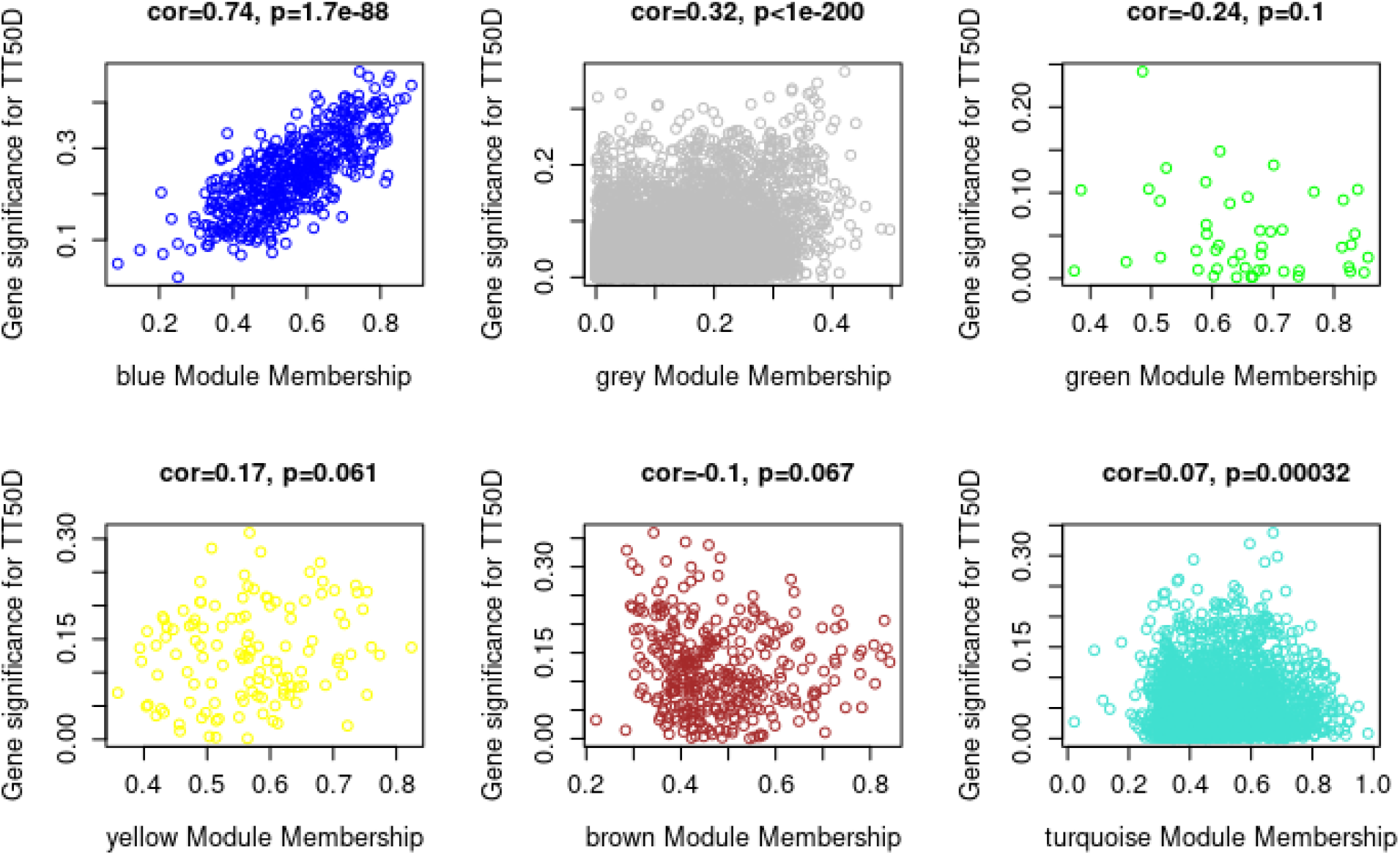
Scatter plots of lncRNA-apoptosis association against module membership. The x-axis represents module membership values and the y-axis represents lncRNA-TT50D correlation values. TT50D: the time in minutes for 50% of cells to become propidium iodide positive post staurosporine-treatment.

A good number of partner protein-coding genes showed highly correlated expression profile with lncRNAs (Supplementary Table VIII). Remarkably, gene ontology analysis of blue module partner protein-coding genes revealed an enrichment in smooth muscle cell relevant terms (Figure 3C and Supplementary Tables IX-XII). This highlights the strength of our proxy approach and suggests that blue module lncRNAs effects on VSMC behaviour may be mediated through their interactions with nearby protein-coding genes.

### Prioritization of candidate blue module gene for experimental validation

To arrive at an lncRNA to characterize from the blue module, we focused those whose expression profile was highly correlated with VSMC behaviour and whose gene module membership value was high. In addition, we also examined their expression profiles on GTEx v8 (GTEx Consortium) and prioritized with high expression in tissues that contain smooth muscle cells. By so doing we selected RP11-400K9.4 (ENSG00000237807) for future functional characterization.

## Discussion

As described earlier, our study discovered 329 lncRNAs that have never been reported in any cell/tissue types. It is well established that the expression levels of IncRNAs are generally lower than protein-coding mRNAs.^2-4^ Their lower abundance makes their more difficult to detect. For lncRNA identification, our study benefited from the removal of rRNAs during RNA library preparation prior to RNA-Seq, as rRNAs are highly abundant in total RNA samples and tend to hinder the detection of other types RNAs in RNA-Seq. Furthermore, the RNA-Seq of our study was performed specifically on VSMCs, rather than using tissue samples comprised of a mix of different types of cells, therefore avoiding potential confounding from other cell types. It is plausible that some or all of the novel lncRNAs identified in our study are unique to VSMCs, raising the possibility that they might have specific roles in VSMCs.

Genetic variants on chromosome 9p21 are associated with several major cardiovascular diseases including coronary artery disease (CAD), abdominal aortic aneurysm and intracranial aneurysm^46, 47^. Although the 9p21 locus has received extensive interests as it was the first locus found to be associated with CAD^46^ and its association with the disease is statistically more significant than any other genetic loci,^48^ the biological mechanism underlying this genetic association is still incompletely understood. Previous studies^49-51^ have shown that the expression of the lncRNA ANRIL (also known as CDKN2B-AS1) is influenced by CAD-associated genetic variants at the 9p21 locus,^46, 48^ and it has been reported that ANRIL can promote migration of VSMCs^42^ and induce apoptosis of a HEK 293 (human embryonic kidney) cell line^52^. In agreement, our study shows that the expression of ANRIL positively correlated with both migration and apoptosis of VSMCs, providing further evidence supporting the role of this lncRNA in modulating VSMC behavior.

## Acknowledgements

This work was supported by grants from the British Heart Foundation (RG/16/13/32609, RG/19/9/34655, SP/19/2/344612, PG/18/73/34059, PG/16/9/31995). The work falls under the portfolio of research conducted within the NIHR Leicester Biomedical Research Centre.

## References

1. ENCODE_Project_Consortium. An integrated encyclopedia of DNA elements in the human genome. Nature. 2012;489:57–74.

2. Djebali S, Davis CA, Merkel A, et al. Landscape of transcription in human cells. Nature. 2012;489:101–108.

3. Hon CC, Ramilowski JA, Harshbarger J, et al. An atlas of human long non-coding rnas with accurate 5’ ends. Nature. 2017;543:199–204.

4. Cabili MN, Trapnell C, Goff L, Koziol M, Tazon-Vega B, Regev A, Rinn JL. Integrative annotation of human large intergenic noncoding rnas reveals global properties and specific subclasses. Genes Dev. 2011;25:1915–1927.

5. Mele M, Ferreira PG, Reverter F, et al. Human genomics. The human transcriptome across tissues and individuals. Science. 2015;348:660–665.

6. Kornienko AE, Dotter CP, Guenzl PM, Gisslinger H, Gisslinger B, Cleary C, Kralovics R, Pauler FM, Barlow DP. Long non-coding rnas display higher natural expression variation than protein-coding genes in healthy humans. Genome Biol. 2016;17:14.

7. Batista PJ, Chang HY. Long noncoding rnas: Cellular address codes in development and disease. Cell. 2013;152:1298–1307.

8. Huang CK, Kafert-Kasting S, Thum T. Preclinical and clinical development of noncoding rna therapeutics for cardiovascular disease. Circ Res. 2020;126:663–678.

9. Jiang MC, Ni JJ, Cui WY, Wang BY, Zhuo W. Emerging roles of lncrna in cancer and therapeutic opportunities. Am J Cancer Res. 2019;9:1354–1366.

10. Hung J, Miscianinov V, Sluimer JC, Newby DE, Baker AH. Targeting non-coding rna in vascular biology and disease. Front Physiol. 2018;9:1655.

11. Uszczynska-Ratajczak B, Lagarde J, Frankish A, Guigo R, Johnson R. Towards a complete map of the human long non-coding rna transcriptome. Nat Rev Genet. 2018;19:535–548.

12. de Goede OM, Nachun DC, Ferraro NM, et al. Population-scale tissue transcriptomics maps long non-coding rnas to complex disease. Cell. 2021;184:2633–2648 e2619.

13. Kim-Hellmuth S, Aguet F, Oliva M, et al. Cell type-specific genetic regulation of gene expression across human tissues. Science. 2020;369:1332-+.

14. Basatemur GL, Jorgensen HF, Clarke MCH, Bennett MR, Mallat Z. Vascular smooth muscle cells in atherosclerosis. Nat Rev Cardiol. 2019;16:727–744.

15. Uchida S, Dimmeler S. Long noncoding rnas in cardiovascular diseases. Circ Res. 2015;116:737–750.

16. Zhang Z, Salisbury D, Sallam T. Long noncoding rnas in atherosclerosis: Jacc review topic of the week. J Am Coll Cardiol. 2018;72:2380–2390.

17. Leik CE, Willey A, Graham MF, Walsh SW. Isolation and culture of arterial smooth muscle cells from human placenta. Hypertension. 2004;43:837–840.

18. Dobin A, Gingeras TR. Mapping rna-seq reads with star. Curr Protoc Bioinformatics. 2015;51:11 14 11–11 14 19.

19. Yates AD, Achuthan P, Akanni W, et al. Ensembl 2020. Nucleic Acids Res. 2020;48:D682–D688.

20. Pertea M, Pertea GM, Antonescu CM, Chang TC, Mendell JT, Salzberg SL. Stringtie enables improved reconstruction of a transcriptome from rna-seq reads. Nat Biotechnol. 2015;33:290–295.

21. Volders PJ, Anckaert J, Verheggen K, Nuytens J, Martens L, Mestdagh P, Vandesompele J. Lncipedia : Towards a reference set of human long non-coding rnas. Nucleic Acids Res. 2019;47:D135–D139.

22. Trapnell C, Roberts A, Goff L, et al. Differential gene and transcript expression analysis of rna-seq experiments with tophat and cufflinks. Nat Protoc. 2012;7:562–578.

23. Pertea G, Pertea M. Gff utilities: Gffread and gffcompare. F1000Res. 2020;9

24. Shen W, Le S, Li Y, Hu F. Seqkit: A cross-platform and ultrafast toolkit for fasta/q file manipulation. PLoS One. 2016;11:e0163962.

25. Kong L, Zhang Y, Ye ZQ, Liu XQ, Zhao SQ, Wei L, Gao G. Cpc: Assess the protein-coding potential of transcripts using sequence features and support vector machine. Nucleic Acids Res. 2007;35:W345–349.

26. Wang L, Park HJ, Dasari S, Wang S, Kocher JP, Li W. Cpat: Coding-potential assessment tool using an alignment-free logistic regression model. Nucleic Acids Res. 2013;41:e74.

27. Li A, Zhang J, Zhou Z. Plek: A tool for predicting long non-coding rnas and messenger rnas based on an improved k-mer scheme. BMC Bioinformatics. 2014;15:311.

28. Wucher V, Legeai F, Hedan B, et al. Feelnc: A tool for long non-coding rna annotation and its application to the dog transcriptome. Nucleic Acids Res. 2017;45:e57.

29. UniProt C. Uniprot: A worldwide hub of protein knowledge. Nucleic Acids Res. 2019;47:D506–D515.

30. Finn RD, Bateman A, Clements J, et al. Pfam: The protein families database. Nucleic Acids Res. 2014;42:D222–230.

31. Camacho C, Coulouris G, Avagyan V, Ma N, Papadopoulos J, Bealer K, Madden TL. Blast+: Architecture and applications. BMC Bioinformatics. 2009;10:421.

32. Bray NL, Pimentel H, Melsted P, Pachter L. Near-optimal probabilistic rna-seq quantification. Nat Biotechnol. 2016;34:525–527.

33. Soneson C, Love MI, Robinson MD. Differential analyses for rna-seq: Transcript-level estimates improve gene-level inferences. F1000Res. 2015;4:1521.

34. Love MI, Huber W, Anders S. Moderated estimation of fold change and dispersion for rna-seq data with deseq2. Genome Biol. 2014;15:550.

35. Drost HG, Paszkowski J. Biomartr: Genomic data retrieval with r. Bioinformatics. 2017;33:1216–1217.

36. Langfelder P, Horvath S. Wgcna: An r package for weighted correlation network analysis. BMC Bioinformatics. 2008;9:559.

37. Liu B, Pjanic M, Wang T, et al. Genetic regulatory mechanisms of smooth muscle cells map to coronary artery disease risk loci. Am J Hum Genet. 2018;103:377–388.

38. Azlan A, Obeidat SM, Yunus MA, Azzam G. Systematic identification and characterization of aedes aegypti long noncoding rnas (lncrnas). Sci Rep. 2019;9:12147.

39. Bush SJ, Muriuki C, McCulloch MEB, Farquhar IL, Clark EL, Hume DA. Cross-species inference of long non-coding rnas greatly expands the ruminant transcriptome. Genet Sel Evol. 2018;50:20.

40. Sarropoulos I, Marin R, Cardoso-Moreira M, Kaessmann H. Developmental dynamics of lncrnas across mammalian organs and species. Nature. 2019;571:510–514.

41. Zhao Q, Sun Y, Wang D, Zhang H, Yu K, Zheng J, Zuo Z. Lncpipe: A nextflow-based pipeline for identification and analysis of long non-coding rnas from rna-seq data. J Genet Genomics. 2018;45:399–401.

42. Zhang C, Ge S, Gong W, Xu J, Guo Z, Liu Z, Gao X, Wei X, Ge S. Lncrna anril acts as a modular scaffold of wdr5 and hdac3 complexes and promotes alteration of the vascular smooth muscle cell phenotype. Cell Death Dis. 2020;11:435.

43. Rinn JL, Chang HY. Genome regulation by long noncoding rnas. Annu Rev Biochem. 2012;81:145–166.

44. Zinad HS, Natasya I, Werner A. Natural antisense transcripts at the interface between host genome and mobile genetic elements. Front Microbiol. 2017;8:2292.

45. Hu YW, Guo FX, Xu YJ, et al. Long noncoding rna nexn-as1 mitigates atherosclerosis by regulating the actin-binding protein nexn. J Clin Invest. 2019;129:1115–1128.

46. Samani NJ, Erdmann J, Hall AS, et al. Genomewide association analysis of coronary artery disease. N Engl J Med. 2007;357:443–453.

47. Helgadottir A, Thorleifsson G, Magnusson KP, et al. The same sequence variant on 9p21 associates with myocardial infarction, abdominal aortic aneurysm and intracranial aneurysm. Nat Genet. 2008;40:217–224.

48. Nelson CP, Goel A, Butterworth AS, et al. Association analyses based on false discovery rate implicate new loci for coronary artery disease. Nat Genet. 2017;49:1385–1391.

49. Jarinova O, Stewart AF, Roberts R, et al. Functional analysis of the chromosome 9p21.3 coronary artery disease risk locus. Arterioscler Thromb Vasc Biol. 2009;29:1671–1677.

50. Holdt LM, Beutner F, Scholz M, Gielen S, Gabel G, Bergert H, Schuler G, Thiery J, Teupser D. Anril expression is associated with atherosclerosis risk at chromosome 9p21. Arterioscler Thromb Vasc Biol. 2010;30:620–627.

51. Motterle A, Pu X, Wood H, et al. Functional analyses of coronary artery disease associated variation on chromosome 9p21 in vascular smooth muscle cells. Hum Mol Genet. 2012;21:4021–4029.

52. Holdt LM, Stahringer A, Sass K, et al. Circular non-coding rna anril modulates ribosomal rna maturation and atherosclerosis in humans. Nat Commun. 2016;7:12429.

53. Zhen-Dong S, Yan H, Zhao-Yue Z, et al. iLoc-lncRNA: predict the subcellular location of lncRNAs by incorporating octamer composition into general PseKNC. Bioinformatics. 2018;34(24): 4196–4204

54. Wang, H., Ding, Y., Tang, J. et al. Identify RNA-associated subcellular localizations based on multi-label learning using Chou’s 5-steps rule. BMC Genomics 22, 56 (2021).

55. Ana Kozomara, Maria Birgaoanu, Sam Griffiths-Jones, miRBase: from microRNA sequences to function, Nucleic Acids Research. 2019;47(1):155–162.

56. Kertesz, M., Iovino, N., Unnerstall, U. et al. The role of site accessibility in microRNA target recognition. Nat Genet. 2007;39:1278–1284

57. van Heesch, S., van Iterson, M., Jacobi, J. et al. Extensive localization of long noncoding RNAs to the cytosol and mono- and polyribosomal complexes. Genome Biol. 2014;15:R6

58. Carlevaro-Fita J, Johnson R. Global positioning system: understanding long noncoding RNAs through subcellular localization. Molecular cell. 2019;73(5):869–83.

59. Aaron R. Quinlan, Ira M. Hall, BEDTools: a flexible suite of utilities for comparing genomic features, Bioinformatics. 2010;26(6):841–842

60. J. Chen, E.E. Bardes, B.J. Aronow, A.G. Jegga ToppGene suite for gene list enrichment analysis and candidate gene prioritization Nucleic Acids Res. 2009;37:W305–11

61. Supek, F., Škunca, N., Repar, J., Vlahoviček, K. and Šmuc, T., 2010. Translational selection is ubiquitous in prokaryotes. PLoS genetics. 2010;6(6):p.e1001004.

62. GTEx Consortium, 2020. The GTEx Consortium atlas of genetic regulatory effects across human tissues. Science, 2020;369(6509):1318–1330.

